# Lineage-specific trait variation generates widespread, contemporaneous coexistence and competitive exclusion dynamics in an invasive, multihost wildlife parasite

**DOI:** 10.1101/2024.11.24.625051

**Authors:** Trenton WJ Garner, Xavier A Harrison, Andy Fenton, Pria N Ghosh, Ruhan Verster, Kris A Murray, Lola M Brookes, Bryony E Allen, Rhys A Farrer, Benedikt R Schmidt, Kieran A Bates, Natasha Kruger, Abigail Wolmarans, Jaime Bosch, Matthew C Fisher, Ché Weldon

## Abstract

Invasive and highly virulent parasites are emerging worldwide, transported to new locations and into novel hosts by anthropogenic activities. Multiple introduction events lead to interactions amongst genetically dissimilar genotypes that can result in either competitive exclusion, coexistence, or cycling through a combination of the two. Here, we report how intra-lineage trait variation of the multihost amphibian parasite *Batrachochytrium dendrobatidis* drives contrasting outcomes of inter-lineage interactions on two continents, Europe and Africa. Through field surveys, competition experiments and mathematical modelling we show that interactions between the same two lineages in the two continents and in different hosts demonstrate both ends of the exclusion/coexistence continuum. Trait variation in one of the two predominating lineages, *Bd*GPL, is responsible for these contrasting outcomes: In Europe, *Bd*GPL is highly competitive and has constrained the distribution of the other, *Bd*CAPE, to two locations. In Africa, *Bd*GPL and *Bd*CAPE can mutually invade host populations when the other lineage is already resident, potentially leading to coinfections and recombination. That these contrasting outcomes are prolonged and contemporaneous for the same two lineages shows that epidemiological models of invasive parasites need to account for trait variation both within and across lineages.

## INTRODUCTION

Introduced, virulent parasites expand their geographic ranges predominantly through host switches (Jones et al. 2008). Co-opting a new host population typically leads to rapid parasite phylogenetic diversification, which in turn can spawn competitive interactions amongst parasite genotypes that differ in traits governing host exploitation (Quigley et al. 2017). The broadly accepted pattern is that less infectious and/or virulent lineages are then replaced with, or are geographically curtailed by, more exploitative variants. The recent SARS-CoV-2 pandemic is one such example, where phylogenetically novel and more infectious lineages globalized by human movements continue to sequentially dominate new cases at regional and international scales (Attwood et al. 2022).

Many wildlife parasites have undergone recent and rapid host and geographic range expansions through human activities, and phylodynamic studies have revealed a contrasting pattern: distributional overlaps amongst diverged genotypes and relative broad host species ranges (Dellicour et al. 2020; O’Hanlon et al. 2019; Streicker et al. 2019; Tracy et al. 2019; Martel et al. 2014; Lawson et al. 2012). Spillovers involving wildlife parasites that can infect multiple host species by definition occupy broader niches than host-specialists. This, in theory, supports parasite phylogenetic diversity without the concordance between phylogeny and phenotype that drive replacement dynamics. Under these circumstances, sequential introductions of divergent lineages in novel host assemblages could result in more widespread lineage coexistence (Boots et al 2014). Replacement dynamics are therefore not a forgone outcome, and for multihost parasites, relaxation of replacement dynamics should be more likely to result in the maintenance of diverged lineages in sympatry. In some viral and bacterial pathogens, genotypes coexist, compete and achieve dominance while still exhibiting evidence of intralineage recombination, suggesting that competitive interactions have multiple solutions (Jackson et al. 2021; Cowley et al. 2018; Bashley 2015; Mehtälä et al. 2013).

If lineage interactions are less predictable in multihost pathogens, the question is then raised if parasites that are highly virulent are consistently competitive. On the face of it, the presence of a competitively dominant parasite should preclude coexistence, but in theory competitive exclusion is only one of several possible outcomes of competition when parasites coinfect hosts (Bashey 2015). While the outcome of competition is largely governed by growth rate, faster is not always better, with the result that more infectious and virulent parasites may be less likely to be transmitted in some circumstances (Althouse & Hanley 2015). Under these circumstances, constraints on traits that favour virulence and competitive ability could be relaxed and the emergence of trait variation that favours competitive and coexistence strategies can result. In support of this, studies have shown how antibiotic-resistant bacteria can spontaneously generate new, functionally diverged phenotypes under competition (Koch et al. 2014). The question then becomes is if highly virulent parasites can maintain multiple competitive strategies that can co-occur in time and space at epidemiologically relevant scales. Again, the expectation in a globalized world is that one or the other strategy should dominate given enough time. However, this remains largely untested. If this were to be the case, though, this would have significant impacts on how wildlife parasite controls are designed and implemented.

*Batrachochytrium dendrobatidis* is a globally widespread and still emerging multihost parasite threatening the existence of hundreds of amphibian species (Scheele et al. 2019). It is generally considered to be the most virulent wildlife pathogen ever recorded (Fisher & Garner 2020). *B. dendrobatidis* emerged from an ancestral Asian centre of genetic diversity presumably through the global trade in amphibians for food, research, pets and ornamentation (O’Hanlon et al. 2019; Wombwell et al. 2016; Fisher & Garner 2007; Garner et al. 2006). Five deeply diverged lineages and several recombinants are currently recognized, and while phylogenies have been used to infer timing and patterns of introduction and diversification of *B. dendrobatidis* lineages (Farrer et al. 2011; O’Hanlon et al. 2018; Byrne et al. 2019; Rothstein et al. 2021), spatiotemporal interactions amongst *Bd* lineages are largely undetermined (but see Jenkinson et al. 2016 and Carvalho et al. 2023 for spatial interactions in Latin America). Current data indicates that for the most part lineages have discrete distributions and rarely appear to interact in both space and time. The Global Pandemic Lineage (*Bd*GPL) is the only lineage that has achieved pan continental distribution and is associated with the great majority of mass mortality events, including locations in Australia, both American continents and most of Europe (Fisher & Garner 2020; O’Hanlon et al. 2018; Byrne et al 2019). Significant contact zones between *Bd*GPL and other lineages have been detected in South America and Africa, and in both locations rare recombinants have been isolated and sequenced (Verster et al. 2024; O’Hanlon et al. 2018; Byrne et al. 2019). In Africa, *Bd*GPL co-occurs with *Bd*CAPE, a lineage that has been detected on three continents and which is largely responsible for severe outbreaks of chytridiomycosis caused by *B. dendrobatidis* that have not been attributed to *Bd*GPL (O’Hanlon et al. 2018). More specifically, mass mortality events and amphibian population declines caused by *Bd*CAPE have been reported in wild amphibians on two of the three continents where *Bd*CAPE occurs, Africa and Europe (Sewell et al. 2021; Weldon et al. 2020; O’Hanlon et al. 2018; Byrne et al 2019; Griffiths et al. 2018; Doddington et al. 2013).

The spatial distributions of *Bd*GPL and *Bd*CAPE on Europe and Africa appear to present somewhat contrasting patterns (Fig. 1a). In Africa, both *Bd*GPL and *Bd*CAPE have been detected in several countries, and it is in one of these countries (Tanzania) where *Bd*CAPE has caused extinction in the wild of the Kihansi spray toad (Sewell et al. 2021; Weldon et al. 2020). Recently, we have described overlapping distributions in South Africa, where we have isolated both lineages from hosts in the same breeding pond, and, in one case, co-infecting individual hosts (Verster et al. 2024; Fig 1a). By contrast, in Europe, *Bd*GPL occurs extensively whereas *Bd*CAPE is restricted to just two geographically distant locations. In this study we sought to determine whether the different distributions within and amongst continents could be attributed to shifts in functional parasite trait values that affect the ability to exploit a host, and whether these concord with phylogenetics. To do this we first evaluated via clustering analysis the currently described distributions of the two lineages in Europe and Africa to highlight their contrasting continental occupancies. We then undertook a series of *in vivo* experiments using European and African amphibian host species to i) experimentally test if infectivity and/or virulence of isolates were attributable to lineage or the continent where we collected them, and; ii) experimentally test if infection dynamics under competition were consistent with patterns of infectivity and virulence measured in the absence of competition. We then used these data to parameterise a multi-lineage epidemiological model to predict whether competitive effects altered the ability of a novel, introduced lineage to establish in the presence of a resident lineage, with implications for lineage co-circulation or exclusion at the host population level. Last, we report what we consider to be direct evidence of lineage replacement in Europe, the continent where we conclude *Bd*GPL has enhanced competitive ability.

**Figure 1.**
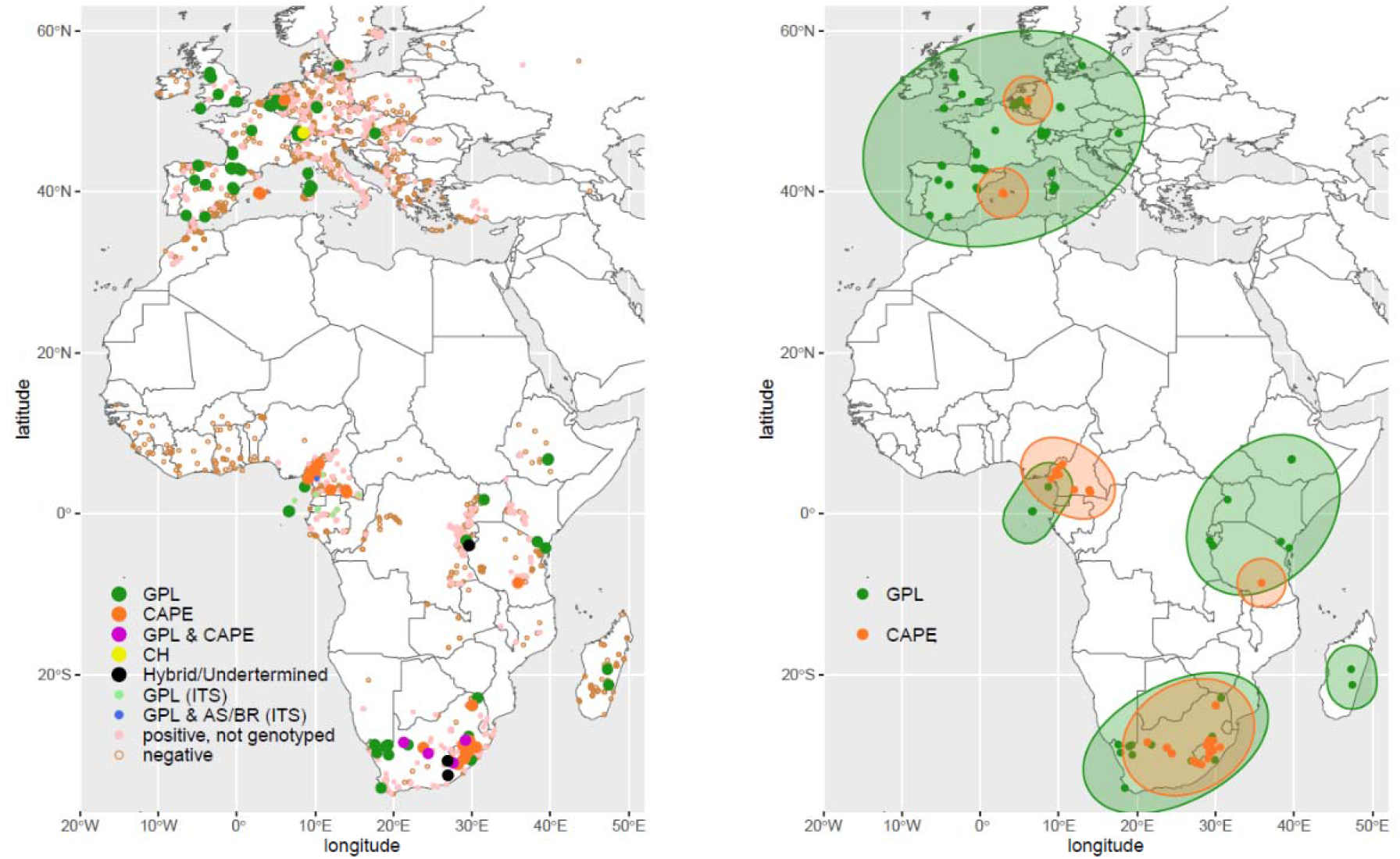
Left panel a) distributions of Bd in Europe and Africa. Large points indicate whole genome sequencing (WGS) lineage genotyping showing exclusive *Bd*CAPE (orange), *Bd*GPL (green), or *Bd*CH (yellow) occurrence. Populations where both lineages were detected are magenta and purported hybrid sites are in black [data from O’Hanlon et al. (2018), Byrne et al. (2019), Sewell et al. (2021), Ghose et al. (2023), and Verster et al. (2024)]. Small points (light green, blue) show putative lineages genotyped in the Gulf of Guinea region via ITS markers only (data from Miller et al. 2018; Hydeman et al. 2017), and therefore currently considered suggestive of genotype (Byrne et al., 2019). Small points (pink) show other Bd-positive records without lineage information, and open circles (brown) show location of negative test results (data from Bd-maps – note most recent updates to 2019 not available for download at time of writing from AmphibianDisease.org, see Koo et al. 2021). Right panel b) Lineage clusters for robust lineage genotypes (BdGPL = green; BdCAPE = orange) across both continents based on single-linkage hierarchical cluster analysis on spatial coordinates only. Note apparently overlapping clusters in Gulf of Guinea region (Central West Africa) are currently distinct with a single-lineage island (BdGPL) and a single-lineage mainland (BdCAPE) clusters.

## METHODS

### Mapping and Cluster Analyses

Mapping data sets were gleaned from multiple sources, including robust data sets involving WGS SNP data from O’Hanlon et al. (2018), multilocus genotyping used by Byrne et al. (2019) and Ghose et al. (2023), lineage-specific qPCR data for *Bd*CAPE and *Bd*GPL published in Verster et al. (2024) and the report of *Bd*CAPE causing species extinction in Tanzania (Sewell et al. 2021). Other records shown in Figure 1a were drawn from publications focussing on distribution and/or lineage diversity of Bd in Africa but where lineage data is either not reported or based on phylogenetically uninformative methods (ITS: see Fig 1 caption for references). Robust lineage records for *Bd*GPL and *Bd*CAPE were subsequently analysed using a simple single-linkage hierarchical cluster approach (*hclust* function in R base stats version 4.3.1) based on purely spatial information (latitude, longitude) using a Great Circle (geographic) distance matrix (*rdist*.*earth* function in the R package ‘fields’). Cluster ellipses were then superimposed over the sites in each cluster (*geom_mark_ellipse* function in the R package ‘ggforce’, which implements the Khachiyan method).

### Infection Experiments Ethics statement

All experimental procedures were subject to ethical review by the Zoological Society of London’s Ethics Committee. Work in the United Kingdom was completed under the conditions of the ‘Animals in Scientific Procedures Act (1986; PPL 80/2466 and PPL P8897246A to Garner, with PIL holders I41A62BDF and I10FA9E48). Field work in South Africa was done under permit to NWU (OP 65/2010).

### Experimental protocol

Animal sources, pre-experiment husbandry and full experimental protocols are available in the supplementary information. More briefly, African and European toadlets (35 per treatment) were exposed 5 (African *Sclerophrys gutturalis*) or 6 (European *Bufo bufo*) times to their respective treatment doses. For both experiments, high and low doses of European *Bd*CAPE were done using isolate TF5a1, isolated from a diseased and moribund *A. muletensis* collected at Torrent de Ferrerets on the island of Mallorca in 2017. High and low doses of European *Bd*GPL were done using IA11 on *B. bufo* and IA042 on *S. gutturalis*, both of which had been isolated from diseased and moribund *Alytes obstetricans* collected from Ibon Acherito in the Spanish Pyrenees (isolated in 2001 and 2017, respectively). High and low doses of South African *Bd*GPL were done using 08MG04 (*B. bufo*) isolated from an apparently healthy *Amietia fuscigula* captured at Silver Mine, KwaZulu-Natal in 2008 and MCT8 (*S. gutturalis*) isolated from a diseased and moribund *Hadromophryne natalensis* sampled in the Royal Natal National Park, KwaZulu-Natal, also in 2008. The South African isolate we used to expose *B. bufo* was SA4C, isolated from an apparently healthy *Amietia angolensis* sampled near Pinetown, KwaZulu-Natal in 2010 (permission for collecting all isolates in O’Hanlon et al. 2018). Date of death or survival to the end of the experiment was recorded and we extracted DNA from an entire hindfoot for qPCR detection and estimation of strength of infection using the standard method of Boyle et al. (2004).

We used parasite-free, singly housed *A. muletensis* tadpoles (*n* = 260, 20 per treatment, 13 treatments in total) to examine opportunities for the competition assay. This is because larval anurans can become infected but death from chytridiomycosis is largely absent until metamorphosis. During early larval stages, infections are restricted to the area surrounding the keratinized mouthparts, meaning that exposures were spatially targeted and co-exposures were far more likely to interact directly (Garner et al. 2009; Blaustein et al. 2005). It also ensured that by swab-sampling tadpole mouthparts we captured the totality of lineage interactions. We used tadpoles of the Mallorcan midwife as the larval stage is protracted, eliminating the possibility of metamorphic death during our experiment (Wells et al. 2015). We exposed tadpoles 8 times to treatments consisting of 4 X early exposures (21000 zoospores in total) followed by 4 X sham exposures, or 8 X exposures (39000 zoospores in total). Negative control animals received 8 X sham exposures. All 4 X exposures followed by sham were done for single lineage exposures, while 8 X exposures were either single lineage exposures, or mixed lineage exposures that alternated lineage order (ie, *Bd*CAPE followed by *Bd*GPL, or vice versa). This design allowed us to compare the effect of lineage competition on post exposure establishment and growth (4 X single lineage exposure against 4 X same followed by competitor), the effect of competition while controlling for dose strength (8 X single lineage versus 8 X coexposures) as well as the effect of dose strength on isolate establishment and growth (4 versus 8). We only competed lineages that came from the same continent (ie African *Bd*GPL against African *Bd*CAPE and European *Bd*GPL against European *Bd*CAPE) and used, respectively 08MG04, SA4C, IA11 and TF5A1. Tadpoles were swabbed before exposures to confirm their infection-free status, and at the experimental endpoint 13 days after the end of exposures. Swabs were then extracted following Boyle et al. (2004) and extraction subjected to lineage-specific qPCRs according to protocols in Ghosh et al. (2021).

### Statistical Analysis of Experimental Data Sets

We conducted all data analysis and visualisation in R v4.2.2 (R Core Team 2022). We fitted models in the Bayesian modelling package *brms* (Bürkner 2018; 2021) to derive means and 95% credible intervals of effect sizes and r^2^ metrics. We used full model tests to assess the importance of models to avoid the inflation of Type I error rate and derive accurate estimates of effect sizes (Forstmeier and Schielzeth 2011). We compared models with the Leave-One-Out (LOO) information criterion and consider the full model to have superior support in the data to the null model when the full model was >6 LOO-IC units lower than the null (Vehtari et al. 2017; 2022).

### Epidemiological Modelling

#### Model structure and assumptions

We developed a mathematical model of lineage co-circulation dynamics within a single host population, illustrated in the schematic shown in Fig. 4A. We assumed one lineage is an endemic ‘resident’ (i.e., it has previously invaded and established to an endemic equilibrium level within the host population) and assessed how the presence of that lineage affected the ability of a novel lineage to invade. For simplicity we modelled infection dynamics within a single ‘season’ of transmission, ignoring host births, and therefore focused on lineage interactions during the initial invasion process of the new lineage.

We assume an initial tadpole population of size *N* at the start of the season, of which a proportion, *p*_*R*_, has been exposed to the resident *Bd* lineage (in what follows the subscript ‘*R*’ refers to the resident lineage); hence the initial number of hosts exposed to the resident lineage is 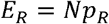, and the remainder are unexposed *(U = N(1* − *p*_*R*_)). The distinction between previously exposed hosts (*E*_*R*_) and unexposed hosts *(U)* allowed us to consider the potential for prior exposure to the resident lineage to alter host susceptibility to, and subsequent infectivity of, a novel invading lineage. Available hosts of classes *U* and *E*_*R*_ are exposed to the invading lineage at a rate dependent on the per capita contact rate (*β*) and the environmental pool of zoospores for that lineage (Z_1_; in what follows the subscript ‘I’ refers to the invading lineage). A proportion, *σ*_*i*_, of these exposed individuals develop patent infections of the invading lineage (i.e., *σ*_*i*_ represents the susceptibility of the host type *i*), where the subscript *i* denotes whether the host was previously unexposed (*i* = *U)* or exposed to the resident (*i*= *R);* if *σ*_*U*_ ≠ *σ*_*R*_, then prior exposure to the resident lineage alters host susceptibility relative to that of unexposed hosts. Hosts that become infected by the invading lineage move to either infected class *I*_*UI*_ or *I*_*RI*_, dependent on whether those hosts had previously been unexposed or exposed to the resident, respectively. Those infected hosts release zoospores into the environmental pool at rates *λ* _*UI*_ and *λ* _*RI*_ respectively; again, these potentially differ to allow for different effects of prior exposure to the resident lineage on shedding rates by the invading lineage. Zoospores are assumed to die at rate *γ*, and we incorporate a low background host mortality rate (*μ*) throughout the season, which is the same for all host classes, regardless of exposure or infection status. The model equations are then:

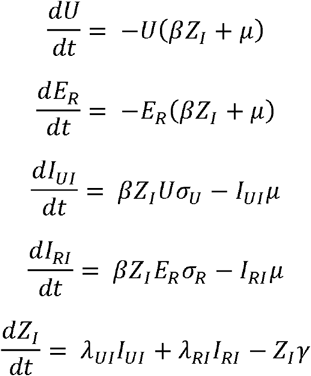

#### Model analysis

*We* used the above framework to calculate the basic reproduction number, *R*_0,*I*_, of the invading lineage, as a measure of its potential to invade and establish within the host population in the presence of the resident lineage. We calculated *R*_0, *I*_ using the next generation approach of Diekmann et al. (2010), where the transmission (*T*) and transition (*G*) matrices, respectively, of the above system of equations, with variables in order {*I*_*UI*_, *I*_*RI*_, *Z*_*I*_}, are:

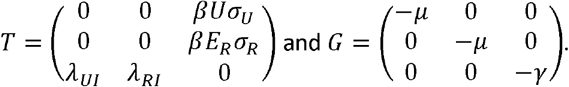

*R*_*0,I*_ is the dominant eigenvalue of the next generation matrix *K = —TG*^-1^ which, since at the point of invasion 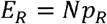 and *U = N(1 — p*_*R*_), is:

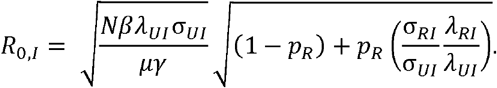

The first square root term is the basic reproduction number of the invading lineage in the *absence* of the resident lineage, whereas the second square root term contains all the co-exposure effects of the resident on the invader. Hence, the change in *R*_*0, I*_ due to the presence of the resident, relative to its baseline value in the absence of the resident, is:

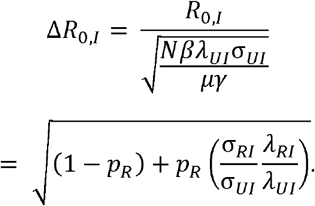

Parameterising this expression using our experimental data (see below), we explored how Δ*R*_0,*I*_ changes across different values of the initial exposure prevalence *p*_*R*_ of the resident lineage. If Δ*R*_0,*I*_ > 1 this indicates that prior circulation of the resident increases the basic reproduction number of the invading lineage, thereby facilitating its likelihood of establishing; if Δ*R*_0,*I*_ < 1 this indicates that the presence of the resident reduces the basic reproduction number of the novel lineage, thereby diminishing its invasion success.

We ran these analyses using the four combinations of both the South African and European pairs of isolates with both *Bd*GPL and BdCAPE taking the roles of resident and invading lineages in turn (i.e., EU *Bd*GPL resident x EU BdCAPE invader; EU BdCAPE resident x EU *Bd*GPL invader. ZA *Bd*GPL resident x ZA *Bd*CAPE invader; ZA *Bd*CAPE resident x ZA *Bd*GPL invader).

#### Model parameterisation

The model parameters to estimate are: (1) the probabilities of infection (the *σ* _i_), and (2) the shedding rates of infected individuals, (the *λ* _*i*_), for each lineage (both European and South African lineages of *Bd*GPL and *Bd*CAPE), and how these might be affected by prior exposure to the other lineage.

#### Probabilities of infection (σ_UI_ and σ _RI_)

Our experiment generated data on the proportion of 20 individuals that became infected with each lineage from each continent under either single or coexposure scenarios; we use these proportions for each combination as estimates of the probabilities of infection (*σ*_*i*_) for each lineage, either alone or when previously exposed to the alternative lineage (Table S1). Note, we restricted analyses to data from our low dose experimental treatments, under the assumption that population-level mean infection loads are likely to be low, particularly during the early stages of an invasion event.

#### Shedding rates (λ_UI;_ and λ_RI_)

Previous work (Allen 2022) has shown a consistent relationship between daily zoospore shedding rate (*λ*, day^−1^) and on-host infection load (*L*) among *A. muletensis* tadpoles, given by: *λ* = 196.17 * *L* (see Supp Info for details). Based on the observed median infection loads from our experiments of the various lineages under the different treatment conditions, we used this relationship to estimate the shedding rates for each lineage under each treatment (Table S1; see Supp Info for additional model details and parameterisation).

#### Field surveys in Switzerland

Methods for aligning lineage-specific mtDNA sequences, candidate primer sequence designs, and assessments of specificity are published elsewhere, along with the conditions for *Bd*GPL and *Bd*CAPE specific Taqman assays (Ghosh et al. 2021). We designed *Bd*CH primer sequences (MTDNACH_F GCGCAGCGAAATCATATAAGATACTT; MTDNACH_R CTCATCGCGGTTGGGTTT) and the MBG FAM-labelled probe sequence (ACTTAAGTATCGAGAACGGTG) following these. Cycle conditions were a modification of Boyle et al. (2004): 50^°^C for two minutes, 95^°^C for ten minutes, followed by 50 cycles including 95^°^C for 15 seconds and 62^°^C for one minute. Detector layer FAM was specified to detect the probe and the cycle threshold was adjusted manually to 0.200 for FAM detector. Standard curves were plotted against a serial dilution of the relevant lineage DNA to generate the linear relationship between Ct and log concentration of GEs. *Bd*CH primers were initially tested against a panel of *Bd*GPL isolates and the one *Bd*CH isolate that was in culture at the time: only this isolate successfully amplified.

Swiss field samples were collected in summer 2011 and April-July 2014 at study sites located in the cantons of Zurich, Baselland and Lucerne, as *Bd*CH was first discovered and described from a pond near the village of Gamlikon, Zurich in 2007: the Gamlikon site was subsequently sampled both in 2011 and 2014 (Table S4). Sites were also selected based on the presence of *Alytes obstetricans*, the species from which *Bd*CH was isolated, using KARCH data (www.karch.ch) and unpublished data (Tobler 2011) to identify *A. obstetricans* breeding sites that were reported as Bd positive. We included a location in Ticino where analysis of museum specimens suggested historical (1901) infection with an unknown lineage (Peyer 2010: Table S4). All field samples were initially tested for infection using the standard Boyle et al. (2004) assay. All samples that tested positive in duplicate for the pan-lineage *Bd* qPCR were reanalyzed using the *Bd*GPL, *Bd*CAPE and *Bd*CH qPCR assays.

## RESULTS

### SPATIAL EVIDENCE SUPPORT CONTINENTAL-SPECIFIC PATTERNS OF LINEAGE COEXISTENCE

In Europe *Bd*GPL formed a single, pan-continental cluster, while in Africa, pan-continental *Bd*GPL formed several clusters and was found on several islands (Fig. 1b). We found at least two distinct multisite clusters of *Bd*CAPE in Africa, while the two *Bd*CAPE sites in Europe do not form a cluster and appear as point introductions, as is the case for *Bd*CAPE in Tanzania. *Bd*CAPE therefore appears constrained within the larger *Bd*GPL cluster in Europe. This is not the case in Africa, where *Bd*CAPE overlaps significantly with the *Bd*GPL cluster in South Africa and predominates around Cameroon (Fig. 1b). Focused surveillance of the two lineages in South Africa identified four locations where both lineages were co-circulating: two west of the Drakensberg range and near the Orange River, a third west of Lesotho but south of (and unconnected to) the Orange River drainage, and a fourth east of Lesotho but on the northern side of the Drakensberg range (Verster et al. 2024). Co-circulation of both lineages was only detected at one location in Europe, in the Netherlands, where *Bd*CAPE was isolated from introduced North American bullfrogs that were first detected in 2010 (Goverse et al. 2012). *Bd*CAPE in Europe was detected elsewhere only on the island of Mallorca, where parasite invasion has been attributed to the reintroduction of Mallorcan midwives that started in 1989: *Bd*GPL has not been detected on the island (Walker et al. 2008; Griffiths et al. 2008, Doddington et al. 2013). The fact that both occurrences are associated with very recent host introductions support the conclusion that invasion is attributable to two recent introduction events that have not spread in either hosts or range.

### EXPERIMENTAL EVIDENCE FOR CONTINENT-SPECIFIC *Bd*GPL INFECTIVITY, VIRULENCE AND COMPETITIVE ADVANTAGE

When exposed to isolates from both continents, European toads died, and with weaker infections, than did African toads, however differences in infectivity and virulence in both species were driven by dose strength and trait divergence in *Bd*GPL. Bayesian linear models revealed support for additive effects of dose and lineage for *B. bufo* (ΔLOO-IC =50.1; Table S1A,B) and a dose:lineage interaction for *S. gutturalis* (ΔLOO-IC =103; Table S1C,D) infection loads. Stronger infections in both toad species occurred when animals were exposed to denser concentrations of European *Bd*GPL zoospores (but not South African *Bd*GPL, Fig. 2A), which in turn generated significantly increased mortality rates (Fig. 2B; Table S1E,F).

**Figure 2.**
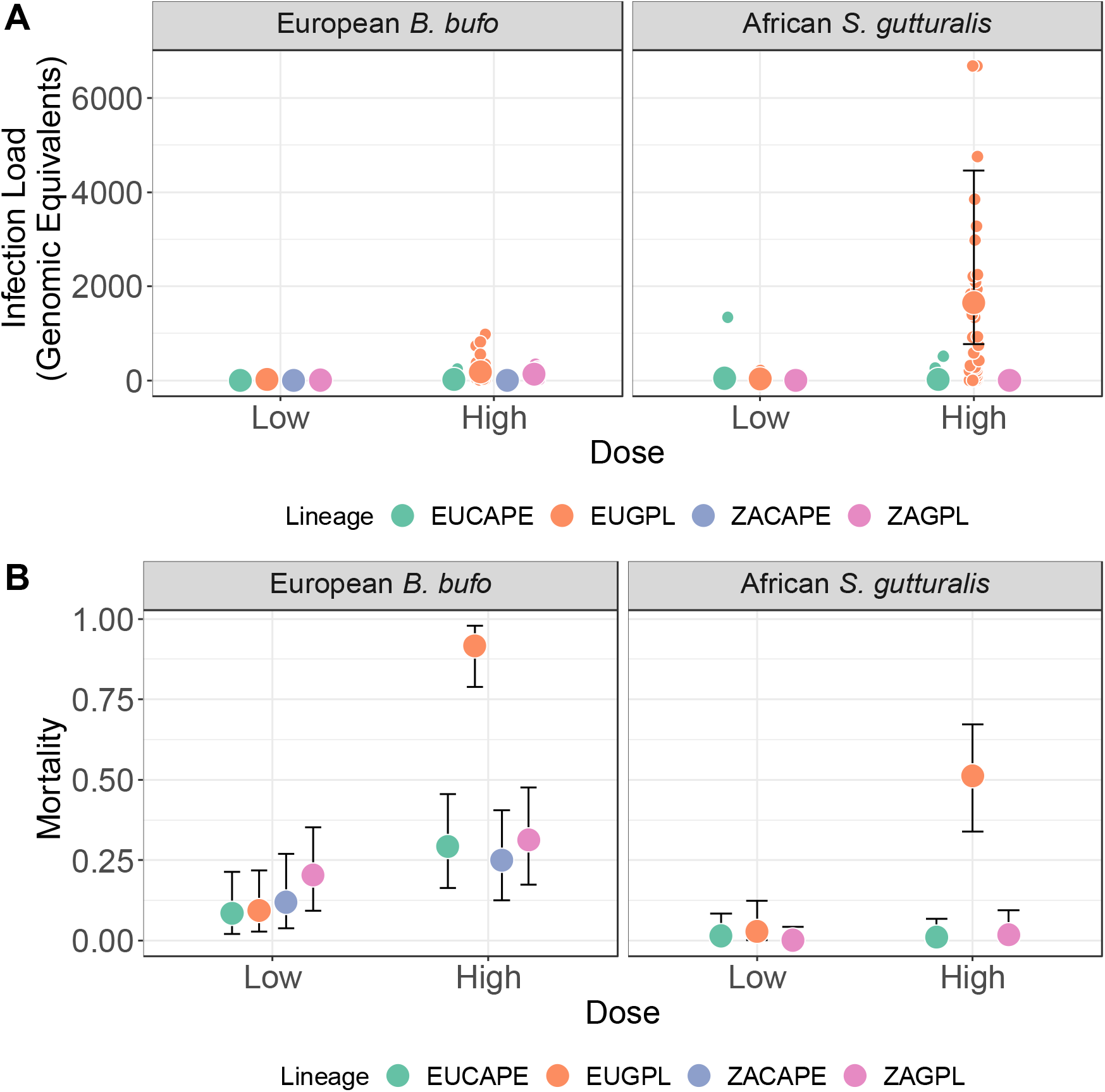
Infection loads (A, top panels) and mortality rates (B, bottom panels) in African Guttural Toads (left panels) and European Common Toads (right panels) exposed to low and high doses of African and European *Bd*GPLs and *Bd*CAPEs. Data missing for African *Bd*CAPE in the Guttural Toad experiment due to isolate unavailability at the time of the experiment. Large points are posterior models means from a Negative Binomial Bayesian linear model, and whiskers are 95% credible intervals. Raw infection data shown as jittered smaller points.

Using Bayesian linear models to explore infection dynamics in tadpoles, we found support for an interaction between the effects of lineage and continent (ΔLOO-IC 19.3; Table S2A) on infection intensities for *Bd*GPL. High dose EU *Bd*GPL treatments generated stronger infections than high dose ZA *Bd*GPL, but we also observed stronger infections of EU *Bd*GPL in the EU *Bd*GPL X EU *Bd*CAPE treatment (Fig. 3A, Fig. S1; Table S2A). The interaction model explained 19.7% of variation in GPL infection intensities (95% credible interval 11.8 – 30.3%). In contrast, we found no evidence of a main effect or interaction in *Bd*CAPE data (ΔLOO-IC = -3.1; Table S2B), where infection loads were consistently low and relatively invariant across all treatments (Fig. 3B). Coinfections were rare (*n* = 3) and restricted to the treatment where exposure order was EU *Bd*CAPE followed by EU *Bd*GPL.

**Figure 3.**
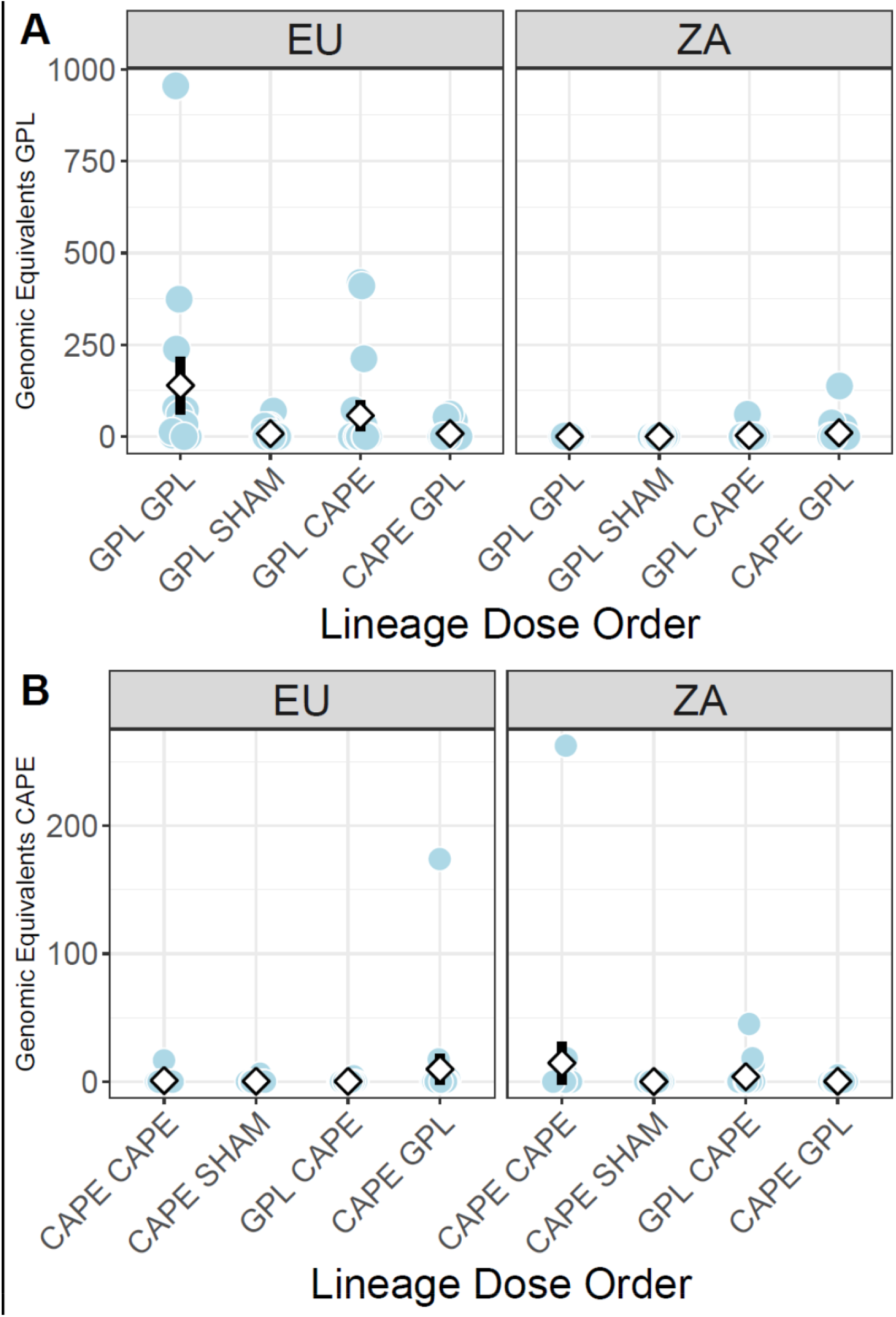
Infection loads (Genomic Equivalents, data scale), split by lineage (A: GPL top, B: CAPE bottom) and by treatment. Diamonds are mean values and bars are S.E. Points have been jittered for display purposes.

**Figure 4.**
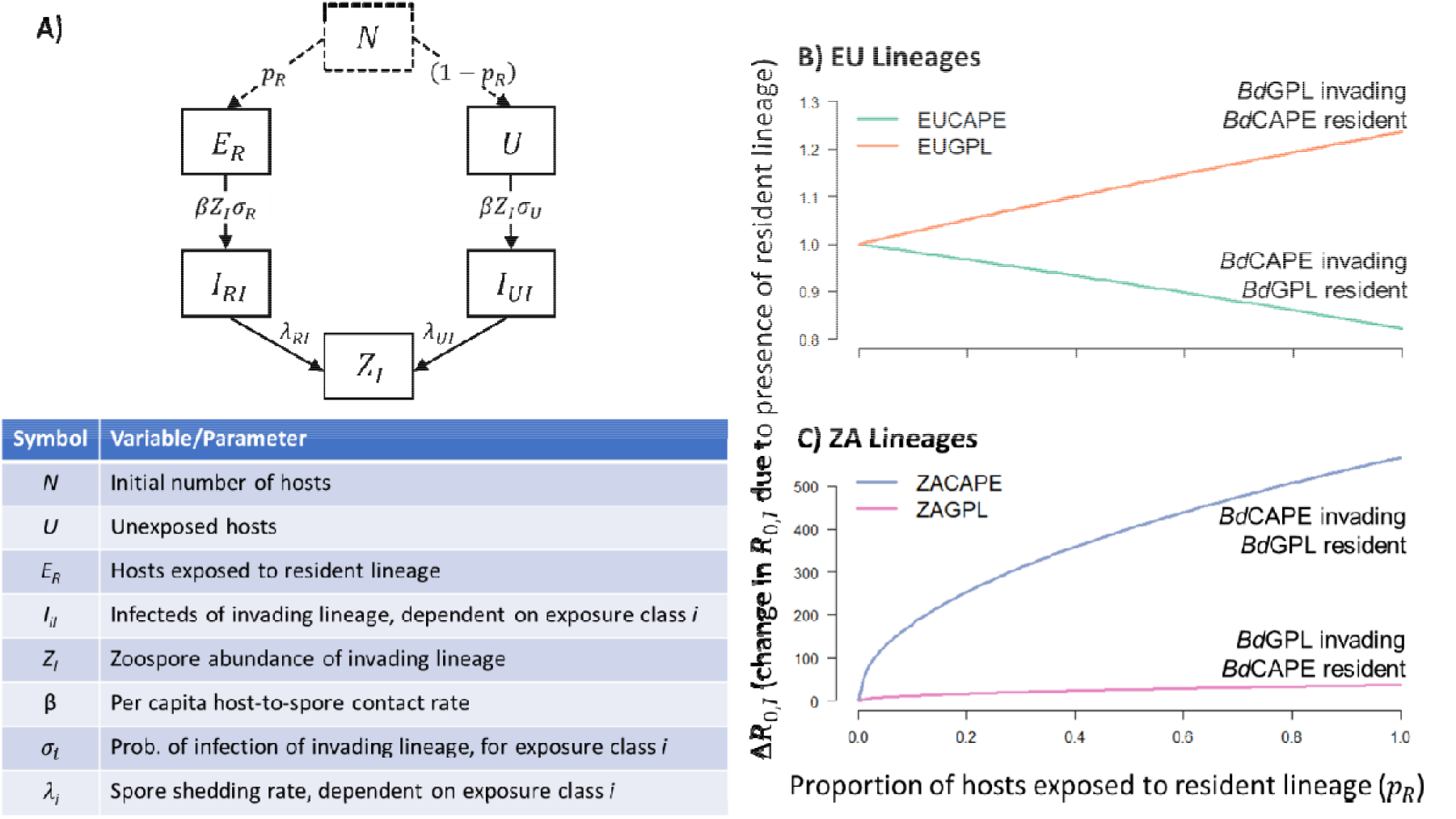
Multi-lineage epidemiological model, predicting the impact of a resident *Bd* lineage on the basic reproduction number of a novel invading lineage. (A) schematic of model structure and parameter definitions. (B) and (C) show changes in the basic reproduction number of the invading *Bd* lineage in the presence of a resident lineage, relative to its basic reproduction number in the absence of the resident (), as a function of the proportion of hosts exposed to the resident lineage (); (B) parameterised for EU lineages and (C) parameterised for ZA lineages, from our infection experiments. Green and blue lines denote the potential for *Bd*CAPE invading in the presence of a *Bd*GPL resident; orange and pink lines denote *Bd*GPL invading in the presence of a *Bd*CAPE resident.

### TRANSMISSION DYNAMICS SUPPORT EUROPEAN *Bd*GPL COMPETITIVE ADVANTAGE WHEN ESTABLISHED AND THE ABILITY TO DISPLACE EUROPEAN *Bd*CAPE

Parameterising our model for the two European isolates from our experiments (Fig. 4B), shows that increasing the initial prevalence of EU *Bd*GPL inevitably led to a reduction in the *R*_0_ of an invading EU *Bd*CAPE lineage (Fig. 4B, green line); hence increasing prevalence of EU *Bd*GPL infections reduced the likelihood of EU *Bd*CAPE invading and establishing. As shown in our competition experiments, EU *Bd*GPL reduces the median infection load of EU *Bd*CAPE infections (Table S3), reducing its transmissibility, thereby reducing its ability to establish. Conversely, our model predicts prior circulation of EU *Bd*CAPE would increase EU *Bd*GPL invasibility (Fig. 4B, orange line), as previous exposure to EU *Bd*CAPE increased EU *Bd*GPL infection loads, and hence zoospore shedding by the invading *Bd*GPL (Table S3). Together these population-level predictions support our hypothesis that *Bd*GPL dominates in Europe by reducing the ability of *Bd*CAPE to establish, and potentially itself benefits in the presence of *Bd*CAPE.

In contrast, the presence of either ZA lineage tended to increase *R*_0_ of the other, and these effects increased as the prevalence of either resident lineage (*p*_R_) increased (Fig. 4C); this effect is particularly clear for ZA *Bd*CAPE invading in the presence of the ZA *Bd*GPL resident (Fig. 4C, blue line). Both lineages benefitted from increased median infection loads following prior infection with the other lineage in our experiments, and *Bd*CAPE also benefitted through increased host susceptibility due to prior exposure to *Bd*GPL (Table S3). Together, this suggests a facilitative effect of each ZA lineage on the other, with no suggestion of competitive effects, driving mutual invasibility, and potentially coexistence of both lineages at the host population level.

### DIRECT EVIDENCE OF LINEAGE REPLACEMENT IN EUROPE

In 2005, we isolated a third European lineage, *Bd*CH, from an artificial pond near the village of Gamlikon in Switzerland (Farrer et al. 2011). This isolate formed a sister group with a lineage composed of isolates entirely derived from Asian amphibians, which in turn forms a cluster distinct from both *Bd*GPL and *Bd*CAPE (Farrer et al. 2011; O’Hanlon et al. 2018). We subsequently conducted surveys of amphibians in the area for evidence of infection with this lineage, first in 2011, restricted to the location where *Bd*CH was initially isolated and focussed on the species from which *Bd*CH was initially isolated (*Alytes obstetricans*, the common midwife). We broadened the survey in 2014 to include both species that occupied the type locality for *Bd*CH (*A. obstetricans* and *Ichthyosaura alpestris*, the alpine newt), alpine newts occupying the pond immediately adjacent to the type locality and series of ponds distributed near to the type locality (Table S4; Schmidt et al. 2023). Overall prevalence of *Bd* across 112 samples was 43% (*n* = 48, 95% CI 0.335 – 0.525). Of these, all samples amplified the *Bd*GPL-specific qPCR assay while all attempts with *Bd*CH-specific or *Bd*CAPE-specific assays were negative (Ghosh et al. 2021).

## DISCUSSION

Taken together, field surveys completed across 20 years, experiments and our theoretical investigation of lineage interactions support the hypothesis that *Bd*GPL in Europe has increased ability to infect, geographically spread and outcompete other lineages than its counterpart in Africa. As a result, European *Bd*GPL appears capable of spatially restricting (*Bd*CAPE) or replacing (*Bd*CH) other lineages despite the availability of numerous suitable host species. This may go some way towards explaining why in Europe host trends appear to be diverging from the global pattern depicted in Scheele et al. (2019). At the global scale, the rate at which new cases of declines attributable to chytridiomycosis appears to be slowing (Luedtke et al. 2023). In Europe, however, spatiotemporal data indicates that infections are geographically spreading and the severity of impacts of lethal chytridiomycosis is increasing on the continent (Martinez Silvestre et al. 2023; Thumsová et al. 2021). The mechanisms underpinning this appear not to be phylogenetically constrained, as we used two different European *Bd*GPL experimental isolates collected almost 20 years apart to generate congruent patterns across the single isolate and competition experiments. European *Bd*GPL isolates also do not form a cluster in the global phylogeny and it is likely *B. dendrobatidis* has been introduced to the European continent on multiple occasions (O’Hanlon et al. 2018). Certainly, this is the case for *Bd*CAPE, which has been introduced twice, over 1500 kilometres and decades apart.

In contrast, both lineages have widespread distributions on the African continent. In South Africa lineage distributions broadly overlap, a pattern that theoretically involves facilitation which enables sequential lineage invasions, cocirculation of highly diverged genotypes, and coinfections. This has provided opportunities for recombination in South Africa, but facilitation has not caused lineage ranges to overlap comprehensively in this country. Environmental envelopes of the two lineages in South Africa are diverged and may be limiting opportunities for the two lineages to interact (Verster et al. 2024). Indeed, the distribution and impacts of *Batrachochytrium dendrobatidis* have been linked to environmental variation since the parasite was first described (Berger et al. 1998; Longcore et al. 1999; Pounds et al. 2006). However, Verster et al. (2024) report unusually frequent lineage co-occurrence along the Orange River in the Northern Cape. Here the climate is significantly warmer and drier than in the best-fit *Bd*CAPE envelope located on and near to the slopes of the part of the Drakensberg Mountains that encapsulate Lesotho. In our model facilitation was strongest when *Bd*CAPE was invading an amphibian population where *Bd*GPL was resident. Lineage cocirculation in the Northern Cape occurred at locations closest to the Free State, one of three provinces within which the majority of proposed *Bd*CAPE envelope occurs (Verster et al. 2024). It is conceivable that downstream migration from the Drakensberg region redistributed *Bd*CAPE to the Northern Cape, where the presence and facilitation of *Bd*GPL enabled persistence in what would otherwise be an inhospitable environment for the lineage. It remains uncertain if South African recombinants exhibit elevated virulence like their South American counterparts might (Greenspan et al. 2018 but see Carvalho et al. 2023). What is clear, however, is that South African recombinants are rare and currently are not significantly expanding their ranges detectably in the regions we sampled. If, as we propose, facilitation of cocirculation is occurring amongst lineages in the Northern Cape, it appears not to facilitate recombination at a similar scale.

What is arguably most striking about our findings is that *B. dendrobatidis* is persistently and simultaneously adopting strategies that occupy either end of the coexistence/competition spectrum. This is a novel finding for a wildlife pathogen, where the evidence base for exploring lineage interactions is largely lacking. Furthermore, these strategies are separated by the slightest of continental divides and are in no way epidemiologically isolated from each other. The trade in exotic amphibians into Europe is largely unregulated, enormous, and highly dynamic in terms of species composition: currently the direction of traffic is into Europe, rather than into Africa (Auliya et al. 2016; Tapley et al. 2011). It is plausible that this dynamic is what sustains the pattern, as the purported source for introductions into Europe would carry genotypes that are not geared towards competitive interactions. Worryingly, the recent emergence of lethal chytridiomycosis in Morocco in North African species that share evolutionary histories with European congeners could presage a reversal of transmission dynamics (Thumsová et al. 2022). Although no invasive amphibians have been recorded, Morocco’s freshwater systems are one of the most highly invaded in Africa (Taybi et al. 2023). Amphibians can be difficult to detect in surveys and early stages of invasions are typically overlooked (Mohanty & Measey 2018). We would argue that surveys for invasive amphibians in this region is a conservation priority. Equally, sequencing of the Moroccan *B. dendrobatidis* is also an urgent matter, to ascertain if these recent North African mortality events are another example of lethal infections due to African *Bd*CAPE or are instead the first records of mortality due to *Bd*GPL on the continent.

The implications of our study are significant for the global effort to conserve amphibian biodiversity. Amphibians are the most threatened vertebrate class and have been since they were first comprehensively assessed in 2004 (Stuart et al. 2004, Luedtke et al. 2023). Chytridiomycosis is responsible for an enormous swather of declines and unusually for a parasite, *B. dendrobatidis* has driven hosts to extinction on more than one continent (Scheele et al. 2019). The effort to control chytridiomycosis in the wild is global, and while the evidence basis for the disease’s role in trimming the amphibian tree of life is vast, control efforts lag far behind. To our knowledge there have been three successful attempts at control since the parasite was first described over 25 years ago, the first of which involved chemical elimination of the parasite in a small landscape (Bosch et al. 2015). This method has proven to be effective at managing outbreaks at amphibian breeding sites at other locations, but scaling up disinfection protocols to manage a globally distributed parasite is problematic (Thumsová et al. 2024). The other two strategies have involved small geographical scale manipulation of the environment to favour host responses and reintroductions of purportedly resistant genotypes into areas where declines have occurred and where the parasite remains in a reservoir species (Waddle et al. 2024; Knapp et al. 2024). Both achievements occurred in a landscape of *Bd*GPL and therefore recorded responses to trait variation within a single genetic lineage of *Batrachochytrium dendrobatidis*. While a previous study illustrated how intralineage competition dynamics may favour parasite tolerance, our results show how interlineage competition dynamics can drive increased virulence in competitive *Bd*GPL genotypes and as well facilitate multilineage interactions that can generate novel, recombined genotypes of unknown pathogenicity (Greener et al. 2020). Multilineage interactions are therefore likely to generate novel infection and disease dynamics in hosts where single lineage responses may not enable tolerance or resistance. For example, amphibian host microbiomes are being extensively explored as a means of mitigating threatening disease dynamics (Walke & Belden 2016). However, even when exposed to single isolates, the capacity of bacterial components of the typical amphibian skin microbiome to inhibit *B. dendrobatidis* growth *in vitro* varied strongly across the *Bd*GPL lineage (Antwis et al. 2015). Given that we have experimentally described increased growth and virulence of European GPL under competition with *Bd*CAPE, we would expect that microbiome profiles that are effective against some *Bd*GPLs in isolation would exhibit decreased efficacy if *Bd*CAPE were to invade the protected population.

The theoretical and experimental approach that we described here has relevance beyond the *Bd*-amphibian system, as animal fungal parasites showing lineage diversity and broad host niches are emerging worldwide. Assumptions of universal outcomes are likely similarly untenable for other globally emerging fungal infections. A case in point is *Candida auris*, first detected in humans in Korea in 1996 then contemporaneously detected on four separate continents (Spivak & Hanson 2018). The five known lineages of *C. auris* are now exhibiting increasingly mixed distributions as human movement erodes previously allopatric lineage distributions; four out of five lineages are now widely invading the USA (Lyman et al. 2023). Simultaneously, this mycosis is showing spillover into other species such as feral dog populations in India and exhibits within and between-lineage trait variation (Yadav et al. 2023; Chybowska et al 2020; Du et al. 2020). The pattern of lineage diversity affecting multiple host species, lineage cocirculation and phenotypic plasticity in traits driving virulence is not uncommon in pathogenic fungi. Accordingly, epidemiological models of emerging fungal parasites should start to account for the potential effects of lineage interactions and dynamic trait variation both within and across lineages.

## Acknowledgements

TWJG, AF, PNG, LAB, BEA, RAF, KAB and MCF were all supported by NERC during their work on this project (NERC standard grants NE/N009967/1, NE/K012509/1 to TWJG, NE/N009800/1 to AF, NE/K014455/1 to MCF, NERC studentships to PNG, BEA, RAF and KAB). TWJG also acknowledges Research England for their ongoing support of the Institute of Zoology.

## References

Allen BE (2022) Transmission of amphibian parasites: exploring the influences of host identity and exposure scenario on key transitions in the transmission pathway. PhD thesis, University of Liverpool

Althouse BM, Hanley KA (2015) The tortoise or the hare? Impacts of within-host dynamics on transmission success of arthropod-borne viruses. Philosophical Transactions of the Royal Society, Series B 370, 20140299

Antwis RE, Preziosi RF, Harrison XA, Garner TWJ (2015) Amphibian symbiotic bacteria do not show a universal ability to inhibit growth of the global panzootic lineage of Batrachochytrium dendrobatidis. Applied and Environmental Microbiology, 81, 3706–3711

Attwood SW, Hill SC, Aanensen DM, Connor TR, Pybus OG (2022) Phylogenetic and phylodynamic approaches to understanding and combating the early SARS-CoV-2 pandemic. Nature Reviews Genetics 23, 547–562

Auliya M, Garćia-Moreno J, Schmidt BR, Schmeller DS, Hoogmoed MS, Fisher MC, Pasmans F, Henle K, Bickford D, Martel A (2016) The global amphibian trade flows through Europe: the need for enforcing and improving legislation. Biodiversity and Conservation 25, 2581–2595

Bashey F (2015) Within-host competitive interactions as a mechanism for the maintenance of parasite diversity. Philosophical Transactions of the Royal Society, Series B 370, 20140301

Berger L, Speare R, Daszak P, Green DE, Cunningham AA, Goggin CL, Slocombe R, Ragan MA, Hyatt AD, McDonald KR, Hines HB, Lips KR, Marantelli G, Parkes H (1998) Chytridiomycosis causes amphibian mortality associated with population declines in the rain forests of Australia and Central America. Proceedings of the National Academy of Sciences of the USA 95, 9031–9036

Blaustein AR, Romansic JM, Scheessele EA, Han BA, Pessier AP, Longcore JE (2005) Interspecific variation in susceptibility of frog tadpoles to the parasiteic fungus Batrachochytrium dendrobatidis. Conservation Biology 19, 1460–1468

Boots M, White A, Best A, Bowers R. 2014 How specificity and epidemiology drive the coevolution of static trait diversity in hosts and parasites. Evolution 68, 1594–1606

Bosch J, Sanchez-Tomé E, Fernández-Lora A, Oliver JA, Fisher MC, Garner TWJ (2015) Successful elimination of a lethal wildlife infectious disease in nature. Biology Letters 11, 20150874

Boyle DG, Boyle DB, Olsen V, Morgan JAT, Hyatt AD (2004) Rapid quantitative detection of chytridiomycosis (Batrachochytrium dendrobatidis) in amphibian samples using real-time Taqman PCR assay. Diseases of Aquatic Organisms 60:141–148

Bürkner P (2018). Advanced Bayesian Multilevel Modeling with the R Package brms. The R Journal, 10(1), 395–411

Bürkner P (2021). Bayesian Item Response Modeling in R with brms and Stan. Journal of Statistical Software, 100(5), 1–54

Byrne AQ, Vredenburg VT, Martel A, Pasmans F, Bell RC, Blackburn DC, Bletz MC, Bosch J, Briggs CJ, Brown RM, Catenazzi A, López MF, Figueroa-Valenzuela R, Ghose SL, Jaeger JR, Jani AJ, Jirku M, Knapp RA, Muñoz A, Portik DM, Richards-Zawacki CL, Rockney H, Rovito SM, Stark T, Sulaeman H, Tao NT, Voyles J, Waddle AW, Yuan Z, Rosenblum EB (2019) Cryptic diversity of a widespread global parasite reveals expanded threats to amphibian conservation. Proceedings of the National Academy of Sciences of the USA 116, 20382–20387

Carvalho T, Medina D, Ribeiro LP, Rodriguez D, Jenkinson TS, Becker CG, Toledo LF, Hite JL (2023) Coinfection with chytrid genotypes drives divergent infection dynamics reflecting regional distribution patterns. Communications Biology 6, 941

Chybowska AD, Childers DS, Farrer R A (2020) Nine Things Genomics Can Tell Us About Candida auris. Frontiers in Genetics 11, 351

Cowley LA, Petersen FC, Junges R, Jimson D. Jimenez M, Morrison DA, Hanage WP (2018) Evolution via recombination: Cell-to-cell contact facilitates larger recombination events in Streptococcus pneumoniae. PLoS Genetics 14, e1007410

Dellicour S, Lequime S, Vrancken B, Gill MS, Bastide P, Gangavarapu K, Matteson NL, Tan Y, du Plessis L, Fisher AA, Nelson MI, Gilbert M, Suchard MA, Andersen KG, Grubaugh ND, Pybus OG, Lemey P (2020) Epidemiological hypothesis testing using a phylogeographic and phylodynamic framework. Nature Communications 11, 5620

Diekmann O, Heesterbeek JAP, Roberts MG (2010) The construction of next-generation matrices for compartmental epidemic models. Journal of the Royal Society Interface 7, 873–885.

Doddington BJ, Bosch J, Oliver JA, Grassly NC, Garcia G, Garner TWJ, Fisher MC (2013) Context-dependent amphibian host population response to an invading parasite. Ecology 94, 1795–1804

Du H, Bing J, Hu T, Ennis CL, Nobile CJ, Huang G (2020) Candida auris: epidemiology, biology, antifungal resistance, and virulence. PLoS Pathogens 16, e1008921

Farrer RA, Weinert LA, Bielby J, Garner TWJ, Balloux F, Clare F, Bosch J, Cunningham AA, Weldon C, du Preez LH, Anderson L, Kosakovsky Pond SL, Shahar-Golan R, Henk DA, Fisher MC (2011) Multiple emergences of amphibian chytridiomycosis include a globalised hypervirulent recombinant lineage. Proceedings of the National Academy of Sciences of the U.S.A. 108, 18732–18736

Fisher MC, Garner TWJ (2020) Chytrid fungi and global amphibian declines. Nature Reviews Microbiology, 18, 332–343

Fisher MC, Garner TWJ (2007) The relationship between the introduction of Batrachochytrium dendrobatidis, the international trade in amphibians and introduced amphibian species. Fungal Biology Reviews 21, 2–9

Forstmeier W, Schielzeth H (2011) Cryptic multiple hypotheses testing in linear models: overestimated effect sizes and the winner’s curse. Behavioral Ecology and Sociobiology 65, 47–55

Garner TWJ, Perkins M, Govindarajulu P, Seglie D, Walker SJ, Cunningham AA, Fisher MC (2006) The emerging amphibian parasite Batrachochytrium dendrobatidis globally infects introduced populations of the North American bullfrog, Rana catesbeiana. Biology Letters 2, 455–459

Garner TWJ, Walker S, Bosch J, Leech S, Rowcliffe JM, Cunningham AA, Fisher MC (2009) Life history trade-offs influence mortality associated with the amphibian parasite Batrachochytrium dendrobatidis. Oikos 118, 783–791

Ghose SL, Yap TA, Byrne AQ, Hasan S, Rosenblum EB, Chan-Alvarado A, CHaukulkar S, Greenbaum E, Koo MS, Kouete MT, Lutz K, McAloose D, Moyer AJ, Parra E, Portik DM, Rockney H, Zink AG, Blackburn DC, Vredenburg VT (2023) Continent-wide recent emergence of a global pathogen in African amphibians. Frontiers in Conservation Science 4

Ghosh P, Verster R, Sewell T, O’Hanlon S, Brookes L, Rieux A, Garner TWJ, Weldon C, Fisher MC (2021) Discriminating lineages of Batrachochytrium dendrobatidis using quantitative PCR. Molecular Ecology Resources, 21, 1452–1459

Goverse E, Creemers R, Spitzen-Van der Sluijs A (2012) Case study on the removal of the American bullfrog in Baarlo, the Netherlands. Report of RAVON (Reptile, Amphibian and Fish Conservation the Netherlands)

Greener MS, Verbrugghe E, Kelly M, Blooi M, Beukema W, Canessa S, Carranza S, Croubels S, De Troyer N, Fernandez-Giberteau D, Goethals P, Lens L, Li Z, Stegen G, Strubbe D, van Leeuwenberg R, Van Praet S, Vila-Escale M, Vervaeke M, Pasmans F, Martel A (2020) Presence of low virulence chytrid fungi could protect European amphibians from more deadly strains. Nature Communications, 11: 5393

Greenspan SE, Lambertini C,Carvalho T, James TY, Toledo LF, Haddad CFB,4 Becker CG (2018) Hybrids of amphibian chytrid show high virulence in native hosts. Scientific Reports 8, 9600

Griffiths RA, Garcia G, Oliver J (2008) Reintroduction of the Mallorcan midwife toad, Mallorca, Spain. In: Global Re-introduction Perspectives: Re-introduction Case-Studies From Around the Globe. Pp. 54, Pritpal S. Soorae, ed. IUCN/SSC Re-introduction Specialist Group, Abu Dhabi, UAE

Griffiths SM, Harrison XA, Weldon C, Wood MD, Pretorius A, Hopkins K, Fix G, Preziosi RF, Antwis RE (2018) Genetic variability and ontogeny predict microbiome structure in a disease-challenged montane amphibian. The ISME Journal 12, 2506–2517

Hydeman ME, Longo AV, Velo-Antón G, Rodriguez D, Zamudio KR, Bell RC (2017) Prevalence and genetic diversity of Batrachochytrium dendrobatidis in Central African island and continental amphibian communities. Ecology and Evolution 7(19), 7729–7738

Jackson B, Boni MF, Bull MJ, Colleran A, Colquhoun RM, Darby AC, Haldenby S, Hill V, Lucaci A, McCrone JT, Nicholls SM, O’Toole Á, Pacchiarini N, Poplawski R, Scher E, Todd F, Webster HJ, Whitehead M, Wierzbicki C, Loman NJ, Connor TR, Robertson DL, Pybus OG, Rambaut A (2021) Generation and transmission of interlineage recombinants in the SARS-CoV-2 pandemic. Cell 184, p5179-5188.e8

Jenkinson TS, Betancourt Román CM, Lambertini C, Valencia-Aguilar A, Rodriguez D, Nunes-de-Almeida CHL, Ruggeri J, Belasen AM, da Silva Leite D, Zamudio KR, Longcore JE, Toledo FL, James TY (2016) Amphibian-killing chytrid in Brazil comprises both locally endemic and globally expanding populations. Molecular Ecology 25, 2978–2996

Jones KE, Patel NG, Levy MA, Storeygard A, Balk D, Gittleman JL, Daszak P (2008) Global trends in emerging infectious diseases. Nature 451, 990–993

Knapp RA, Wilber MQ, Joseph MB, Smith TC, Grasso RL (2024) Reintroduction of resistant frogs facilitates landscape-scale recovery in the presence of a lethal fungal disease. Nature Communications 15, 9436

Koch G, Yepes A, Förstner KU, Wermser C, Stengel ST, Modamio J, Ohlsen K, Foster KR, Lopez D (2014) Evolution of resistance to a last-resort antibiotic in Staphylococcus aureus via bacterial competition. Cell 158, 1060–1071

Koo MS, Vredenberg VT, Deck JB, Olson DH, Ronnenberg KL, Wake DB (2021) Tracking, Synthesizing, and Sharing Global Batrachochytrium Data at AmphibianDisease.org. Frontiers in Veterinary Science 8

Lawson B, Robinson RA, Colville KM, Peck KM, Chantrey J, Pennycott TW, Simpson VR, Toms MP, Cunningham AA (2012) The emergence and spread of finch trichomonosis in the British Isles. Philosophical Transactions of the Royal Society B: Biological Sciences 367, 2852–2863

Longcore JE, Pessier AP, Nichols DK (1999) Batrachochytrium dendrobatidis gen. et sp. nov., a chytrid parasiteic to amphibians. Mycology 91, 219–227

Luedtke JA, Chanson J, Neam K, Hobin L, Maciel AO, Catenazzi A, Borzée A, Hamidy A, Aowphol A, Jean A, Sosa-Bartuano Á, Fong AG, de Silva A, Fouquet A, Angulo A, Kidov AA, Muñoz Saravia A, Diesmos, AC, Tominaga A, Shrestha B, Gratwicke B, Tjaturadi B, Martínez Rivera CC, Vásquez Almazán CR, Señaris C, Chandramouli SR, Strüssmann C, Cortez Fernández CF, Azat C, Hoskin CJ, Hilton-Taylor C, Whyte DL, Gower DJ, Olson DH, Cisneros-Heredia DF, Santana DJ, Nagombi E, Najafi-Majd E, Quah ESH, Bolaños F, Xie F, Brusquetti F, Álvarez FS, Andreone F, Glaw F, Castañeda FE, Kraus F, Parra-Olea G, Chaves G, Medina-Rangel GF, González-Durán G, Ortega-Andrade HM, Machado IF, Das I, Dias IR, Urbina-Cardona JN, Crnobrnja-Isailović J, Yang J-H, Jianping J, Wangyal JT, Rowley JJL, Measey J, Vasudevan K, Chan KO, Gururaja KV, Ovaska K, Warr LC, Canseco-Márquez L, Toledo LF, Díaz LM, Khan MMH, Meegaskumbura M, Acevedo ME, Napoli MF, Ponce MA, Vaira M, Lampo M, Yánez-Muñoz MH, Scherz MD, Rödel M-O, Matsui M, Fildor M, Kusrini MD, Ahmed MF, Rais M, Kouamé NG, García N, Gonwouo NL, Burrowes PA, Imbun PY, Wagner P, Kok PJR, Joglar RL, Auguste RJ, Brandão R, Ibáñez R, von May R, Hedges SB, Biju SD, Ganesh SR, Wren S, Das S, Flechas SV, Ashpole SL, Robleto-Hernández SJ, Loader SP, Incháustegui SJ, Garg S, Phimmachak S, Richards SJ, Slimani T, Osborne-Naikatini T, Abreu-Jardim TPF, Condez TH, De Carvalho TR, Cutajar TP, Pierson TW, Nguyen TQ, Kaya U, Yuan Z, Long B, Langhammer P, Stuart SN (2023) Ongoing declines for the world’s amphibians in the face of emerging threats. Nature 622, 308–314

Lyman M, Forsberg K, Sexton DJ, Chow NA, Lockhart SR, Jackson BR, Chiller T (2023) Worsening Spread of Candida auris in the United States, 2019 to 2021. Annals of Internal Medicine 1 76(4), 489–495

Martel A, Blooi M, Adriaensen C, Van Rooij P, Beukema W, Fisher MC, Farrer RA, Schmidt BR, Tobler U, Goka K, Lips KR, Muletz C, Zamudio K, Bosch J, Lötters S, Wombwell E, Garner TWJ, Spitzen-van der Sluijs A, Salvidio S, Ducatelle R, Nishikawa K, Nguyen TT, Van Bocxlaer I, Bossuyt F, Pasmans F (2014) Recent introduction of a chytrid fungus endangers Western Palearctic salamanders. Science, 346, 630–631

Martinez Silvestre A, Loras-Ortí F, Garcia-Salmeron A, Pujol-Buxó E, Pérez-Novo I, Maluquer-Margalef J, Poch S, Thumsová B, Bosch J (2023) Introduced Mediterranean painted frogs (Discoglossus pictus) are possible supershedders of the fungus Batrachochytrium dendrobatidis in Catalonia (NE Spain). Amphibia-Reptilia, 51, 1–5

Mehtälä J, Antonio M, Kaltoft MS, O’Brien KL, Auranen K (2013) Competition between Streptococcus pneumoniae strains: implications for vaccine-induced replacement in colonization and disease. Epidemiology 24, 522–529

Miller CA, Taboue GCT, Ekane MMP, Robak M, Sesink Clee PR, Richards-Zawacki C, Fokam EB, Fuashi NA, Anthony NM (2018) Distribution modeling and lineage diversity of the chytrid fungus Batrachochytrium dendrobatidis (Bd) in a central African amphibian hotspot. PLoS ONE 13(6), e0199288

Mohanty NP, Measey J (2018) Reconstructing biological invasions using public surveys: a new approach to retrospectively assess spatio-temporal changes in invasive spread. Biological Invasions 21, 467–480

Mutz P, Rochman ND, Wolf YI, Faure G, Zhang F, Koonin EV (2022) Human pathogenic RNA viruses establish noncompeting lineages by occupying independent niches. Proceedings of the National Academy of Sciences of the U.S.A. 119 (23), e2121335119

O’Hanlon SJ, Rieux A, Farrer RA, Rosa GM, Waldman B, Bataille A, Kosch TA, Murray K, Brankovics B, Fumagalli M, Martin MD, Wales N, Alvarado-Rybak M, Berger L, Böll S, Brookes L, Clare F, Courtois EA, Cunningham AA, Doherty-Bone T, Ghosh P, Gower DJ, Hintz WE, Höglund J, Jenkinson TS, Lin C-F, Laurila A, Loyau A, Martel A, Meurling S, Miaud C, Minting P, Pasmans F, Schmeller D, Schmidt BR, Shelton J, Skerratt LF, Smith F, Soto-Azat C, Spagnoletti M, Tessa G, Toledo LF, Valenzuela-Sánchez A, Verster R, Vörös J, Wierzbicki C, Wombwell E, Zamudio KR, Aanensen DM, James TY, Gilbert MTP, Weldon C, Bosch J, Balloux F, Garner TWJ, Fisher MC (2018) Recent Asian origin of chytrid fungi causing global amphibian declines. Science 360, 621–627

Peyer NF (2010) Historical evidence for the presence of the emerging amphibian parasite Batrachochytrium dendrobatidis in Switzerland. M.Sc Thesis, University of Zurich.

Pounds JA, Bustamante MR, Coloma LA, Consuegra JA, Fogden MPL, Foster PN, La Marca E, Masters KL, Merino-Viteri A, Puschendorf R, Ron SR, Sánchez-Azofeifa GA, Still CJ, Young BE (2006) Widespread amphibian extinctions from epidemic disease driven by global warming. Nature 439, 161–167

Quigley BJZ, Brown SP, Leggett HC, Scanlan PD, Buckling A (2017) Within-host interference competition can prevent invasion of rare parasites. Parasitology 145, 770–774

R Core Team (2022) R: A language and environment for statistical computing. R Foundation for Statistical Computing,Vienna, Austria. URL https://www.R-project.org/.

Rothstein AP, Byrne AQ, Knapp RA, Briggs CJ, Voyles J, Richards-Zawacki CL, Rosenblum EB (2021) Divergent regional evolutionary histories of a devastating global amphibian parasite. Proceedings of the Royal Society B Biological Sciences 288, 20210782

Scheele BC, Pasmans F, Berger L, Skerratt LF, Martel A, Beukema W, Acevedo AA, Burrowes PA, Carvalho T, Catenazzi A, De La Riva I, Fisher MC, Flechas SV, Foster CN, Frías-Álvarez P, Garner TWJ, Gratwicke B, Guayasamin JM, Mirschfeld M, Kolby JE, Kosch TA, La Marca E, Lindenmayer DB, Lips KR, Longo AV, Maneyro R, McDonald CA, Mendelson III J, Palacios-Rodriguez P, Parra-Olea G, Richards-Zawacki CL, Rödel M-O, Rovito SM, Soto-Azat C, Toledo LF, Voyles J, Weldon C, Whitfield SM, Milkinson M, Zamudio K, Canessa S (2019) The aftermath of an amphibian fungal panzootic reveals unprecedented loss of biodiversity. Science 363, 1459–1463

Schmidt B, Bohnenstengel T, Zumbach S (2023). Swiss National Amphibian Databank. Swiss National Biodiversity Data and Information Centres – infospecies.ch. Occurrence dataset 10.15468/ggwedn

Sewell TR, van Dorp L, Ghosh PN, Wierzbicki C, Caroe C, Lyakurwa JV, Tonelli E, Bowkett A, Marsden S, Cunningham AA, Garner TWJ, Gilbert TP, Weldon C, Fisher MC (2021) A series of terribly unfortunate events: How environment and infection synergized to cause the Kihansi spray toad extinction. bioRxiv 10.1101/2021.11.10.468118

Spivak ES, Hansen KE (2018) Candida auris: an Emerging Fungal Pathogen. Journal of Clinical Microbiology 56, e01588–17

Stuart SN, Chanson JS, Cox NA, Young BE, Rodrigues ASL, Fischman DL, Waller RW (2004) Status and trends of amphibian declines and extinctions worldwide. Science 306, 1783–6

Streicker DG, González SLF, Luconi G, Barrientos RG, Leon B (2019) Phylodynamics reveals extinction-recolonization dynamics underpin apparently endemic vampire bat rabies in Costa Rica. Proceedings of the Royal Society B Biological Sciences 286, 20191527

Tapley B, Griffiths RA, Bride I (2011) Dynamics of the trade in reptiles and amphibians within the United Kingdom over a ten-year period. Herpetological Journal 21, 27–34

Taybi AF, Mabrouki Y, PIscart C (2023) Distribution of freshwater alien animal species in Morocoo: current knowledge and management issues. Diversity 15, 169

Thumsová, B., Donaire-Barroso, D., El Mouden, E. H. & Bosch, J. (2022). Fatal chytridiomycosis in the Moroccan midwife toad Alytes maurus and potential distribution of Batrachochytrium dendrobatidis across Morocco. African Journal of Herpetology 71, 72–82

Thumsová B, González-Miras E, Faulkner SC, Bosch J (2021) Rapid spread of a virulent amphibian parasite in nature. Biological Invasions 23, 3151–3160

Thumsová B, González-Miras E, Rubio Á, Granados I, Bates KA, Bosch J (2024) Chemical disinfection as a simple and reliable method to control the amphibian chytrid fungus at breeding points of endangered amphibians. Scientific Reports 14, 5151

Tobler U (2011) Differential responses on individual- and population-level to a fungal parasite: Bd infection in the Midwife toad Alytes obstetricans. PhD Thesis, University of Zurich.

Tracy AM, Pielmeier ML, Yoshioka RM, Heron SF, Harvell CD (2019) Increases and decreases in marine disease reports in an era of global change. Proceedings of the Royal Society B Biological Sciences 286, 20191718

Vehtari A, Gelman A, Gabry J (2017). “Practical Bayesian model evaluation using leave-one-out cross-validation and WAIC.” Statistics and Computing, 27, 1413–1432

Vehtari A, Gabry J, Magnusson M, Yao Y, Bürkner P, Paananen T, Gelman A (2022). “loo: Efficient leave-one-out cross-validation and WAIC for Bayesian models.” R package version 2.5.1, https://mc-stan.org/loo/

Verster R, Ghosh PN, Sewell TR, Garner TWJ, Fisher MC, Muller W, Cilliers D, Weldon C (2024) Environment predicts Batrachochytrium dendrobatidis lineage distribution and zones of recombination in South Africa. Ecology and Evolution14, e11037

Waddle AW, Clulow S, Aquilina A, Sauer EL, Kaiser SW, Miller C, Flegg JA, Campbell PT, Gallagher H, Dimovski I, Lambreghts Y, Berger L, Skerratt LF, Shine R (2024) Hotspot shelters stimulate frog resistance to chytridiomycosis. Nature 631, 344–349

Walke JB, Belden LK (2016) Harnessing the microbiome to prevent fungal infections: lessons from amphibians. PLoS Pathogens 12, e1005796

Walker SF, Bosch J, James TY, Litvintseva AP, Valls JAO, Piña S, García G, Rosa GA, Cunningham AA, Hole S, Griffiths R, Fisher MC (2008) Invasive parasites threaten species recovery programs. Current Biology 18, R853–R854

Weldon C, Channing A, Misinzo G, Cunningham AA (2020) Disease driven extinction in the wild of the Kihansi spray toad, Nectophrynoides asperginis. African Journal of Herpetology 69, 151–164

Wells E, Garcia-Alonso D, Rosa GM, García G, Tapley B (2015) Amphibian Taxon Advisory Group Best Practice Guidelines for Midwife toads (Alytes sp.). Version 1.

Wombwell E, Garner TWJ, Cunningham AA, Quest R, Pritchard S, Rowcliffe JM, Griffiths RA (2016) Detection of Batrachochytrium dendrobatidis in amphibians imported into the UK for the pet trade. Ecohealth 13, 456–466

Yadav A, Wang Y, Jain K, Panwar VAR, Kaur H, Kasana V, Xu J, Chowdhary A (2023) Candida auris in dog ears. Journal of Fungi 9, 720

