## Supplementary material for "Lineage-specific trait variation generates widespread, contemporaneous coexistence and competitive exclusion dynamics in an invasive, multihost wildlife parasite": Suppl info

Supplementary information

Research was designed and reported in accordance with the ARRIVE Guidelines. Due to biosecurity risk, blinding was only partially applied (control animals were always identifiable, but specific treatments were concealed). During the laboratory DNA detection work, researcher-blinding was applied by numbering swab samples, which obscured treatment groups. Sample size calculations for experimental designs were based on previous experiments. Sex of study animals remains unknown. All laboratory and animal room waste was either treated with Virkon or autoclaved before incineration.

European and African toad experimental design and implementation

Collection (with landowner permission), pre-experimental husbandry and experiments were conducted in the country to which each species is native. Both *Bufo bufo* and *Sclerophrys gutturalis* were collected as clutches, each kept, hatched and reared as tadpoles in large groups in captivity at ambient temperature and in single outdoor static enclosures (60:40:40 cm) containing aged tapwater (*B. bufo*) or borehole water (*A. gutturalis*). *B. bufo* tadpoles were fed 5 Tetra Tabimin tablets 3 days per week, while *S. gutturalis* tadpoles were fed Tetra Tabimin at a concentration of 1mg/ml enclosure water volume 3 times per week. Once tadpoles had reached Gosner stage 42 or greater, metamorphs were transferred to group housing split between shallow aquatic and terrestrial (sterilized pea gravel) habitats. Cover objects were provided in the terrestrial section. Food was switched to *ad libitum* hatchling crickets dusted with nutrobal (Vetark®) and housing units cleaned twice a week. Once sufficient numbers of metamorphs had been transferred to aquatic/terrestrial housing, metamorphs were then transferred to climate-controlled experimental facilities at the respective institutions, with stocking density set at 40 metamorphs per housing unit. Using a random number generator, experimental animals were randomly assigned to treatments (n=35/treatment) and assigned individual containers. *B. bufo* were housed in 700 mL volume Really Useful Boxes lined with damp (aged tap water), bleach-free paper towelling and with the lid from a small cell culture flask as a cover object. Openings were cut into the side of flask lids to facilitate entry under cover. Experimental units were cleaned and animals fed twice per week. Facilities were kept on a 12:12 light:dark cycle and temperatures kept at 18±2˚C. *A. gutturalis* were housed in 400 mL plastic food pots with lids, with air holes cut into lids. Again, housing was lined with damp (bore hole water) paper towelling, but no cover objects were provided. South African facilities were also kept on a 12:12 light:dark cycle but with temperatures kept at 20±2˚C . For both experiments, experimental units were cleaned and animals fed twice per week.

After two weeks acclimatisation and confirmation of infection-free status via qPCR, *B. bufo* were exposed 6 times over 10 days to a total of 69,000 (high) or 690 zoospores, estimated through direct counts using a haemocytometer, while *S. gutturalis* were exposed 5 times over 20 days to a total of 220,000 (high) or 2,200 (low) zoospores, estimated as above. Metamorphs were exposed individually in petri dishes containing 20 mL of husbandry water, their respective dose (high, low, or negative control composed of chytrid culture media) and for a maximum of 4 hours, then returned to their housing unit. During and after exposures toadlets were fed (approx. n= 6) hatchling and 1^st^ instar crickets at each feed, dusted with nutrobal (Vetark®). Containers were sprayed with husbandry water as needed to maintain damp conditions that were not lethal to feed crickets.

We conducted welfare checks every day, tracked survival daily, and when necessary applied humane euthanasia to animals that exhibited the humane endpoints identified in the UK project license (moribund, lack of righting reflex, inability to feed). Experiments were stopped 31 (*B. bufo*) and 21 (*S. gutturalis*) days after the first exposure, at which point all remaining animals were humanely killed. Humane killing was done using a non-schedule 1 method consisting of an overdose of MS222 followed by immersion in storage ethanol several minutes after the cessation of responsiveness to physical stimulus. Dead and euthanized animals were stored in 70% ethanol until DNA extraction, which was done following the bead-beater method of Boyle et al. (2004). Detection of and estimation of strength of infection was done following method in the same publication, using duplicate amplifications for each animal and known concentration standards (0.1, 1.0. 10 and 100 zsps) for standard curves. Animals were considered infected when both samples amplified and exceeded the 0.1 GE threshold.

Mallorcan midwife tadpoles experimental design and implementation

Tadpoles were excess captive-held zoo amphibians and on arrival to the research facility were housed in climate controlled rooms ~~(~~ambient 18±1C) , under UVB lighting until the start of the experiment and in groups of 40 per tank (in approx. 80l water capacity, really useful boxes) in constantly aerated, aged water. We fed tadpoles4 crushed tablets of fed Tetra Tabimin twice per week with leaf lettuce available throughout). Animals were checked daily for signs of ill-health and animals approaching metamorphosis were removed. Water tests were undertaken weekly, using water from both the stock and experimental housing.

After 1 week acclimatization and confirmation of infection-free status via qPCR, tadpoles were randomly allocated to 700 mL Really Useful Boxes containing 500 mL aged tapwater and randomly assigned to treatments. 260 tadpoles (20 per treatment) were selected for the experiment based on their size and lack of limb buds. Throughout the experiment tadpoles were fed 0.9 ml ground of 1g/100ml Tetra Tabimin suspended in ddH20 every three days, and water was changed completely ever second day during exposures. Tadpoles were exposed every two days for four hours for a total of 8 X [4X isolate followed by sham, or 8 X isolate(s)] exposures. After exposures water changes were reduced to every four days. Tadpole mouthparts were swabbed 13 days after the final exposure, after which the experiment was closed down and all tadpoles humanely killed as above. We monitored tadpole welfare throughout.

Estimating the relationship between infection load and zoospore shedding rate for model parameterisation

Allen (2022) carried out infection experiments on *Alytes muletensis* tadpoles. These tadpoles were bred on site and under license at the Institute of Zoology in climate-controlled rooms (ambient 18±1C). Breeding stocks of *A. muletensis* were kept in large enclosures (1m X 60 cm X 60 cm) at stocking densities not exceeding 20 animals per enclosure and fed crickets dusted with Vetark®) *ad libitum,* 3 days per week. Frogs were left to mate and males to incubate clutches under cover objects undisturbed in enclosures. Hatched tadpoles were collected from large water dishes where males deposit mature clutches. Tadpoles were then reared in groups (max 40 per tank) in constantly aerated 80 litre tanks containing aged water and provided food (4 crushed Tetra Tabimin, twice weekly) and lettuce throughout. Frogs produced clutches over several months to generate sufficient tadpoles. Following the experiment we used q-PCR to quantify both infection load (genomic equivalents) and zoospore shedding (genomic equivalents, equivalent to a single zoospore, over 4 hours, using a modified soak procedure based on DiRenzo et al. (2014) and Reeder et al. (2012)) at 9 days post exposure. Assuming the same shedding rate over 24 hours (i.e., multiplying the observed GEs released over 4h by six) shows a positive relationship between on-host load (*L*) and daily spore shedding rate ($\lambda$) of $\lambda=196.17*L$ (Fig. S2). We use this relationship to convert the observed median infection loads from our experiments to estimate daily shedding rates of the different lineages under the different treatment scenarios (Table S1).

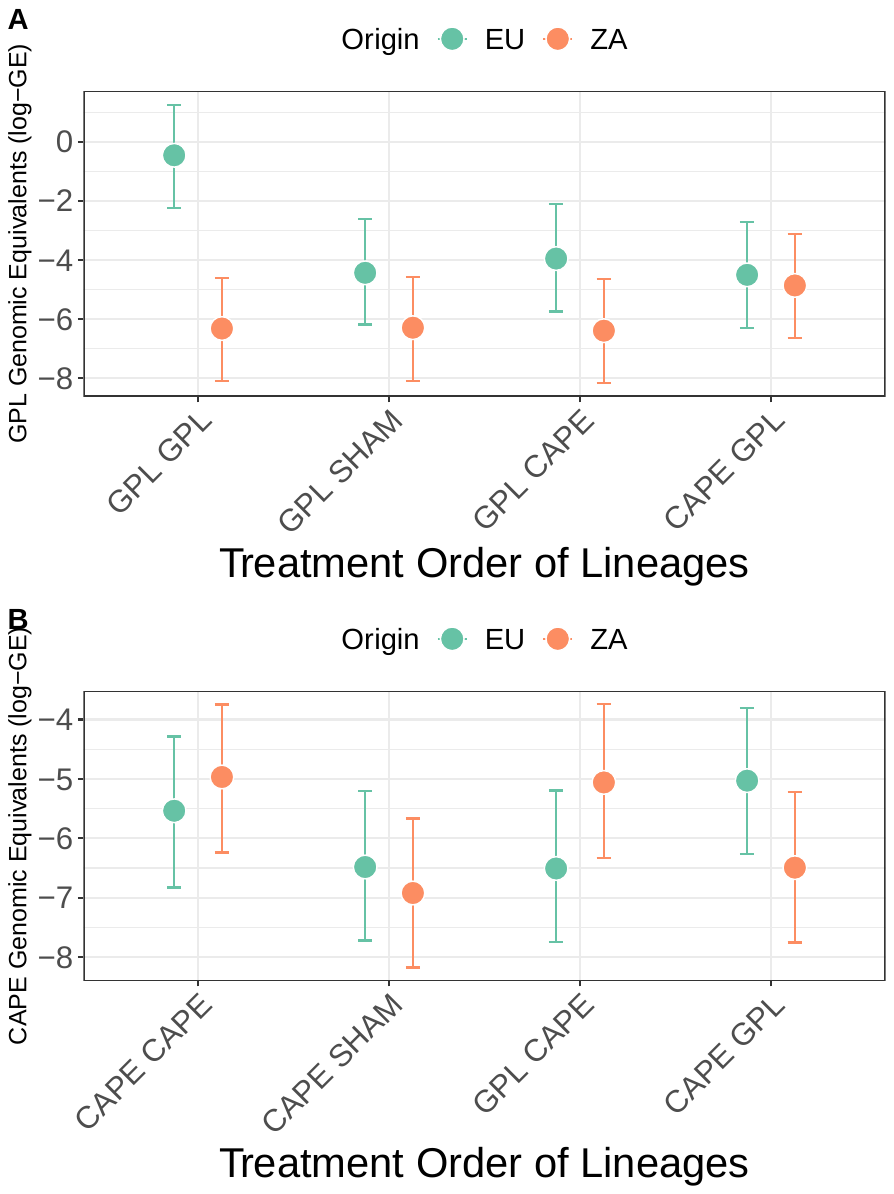

Figure S1. Model predictions of infection intensities (log scale) for treatment order and lineage effects

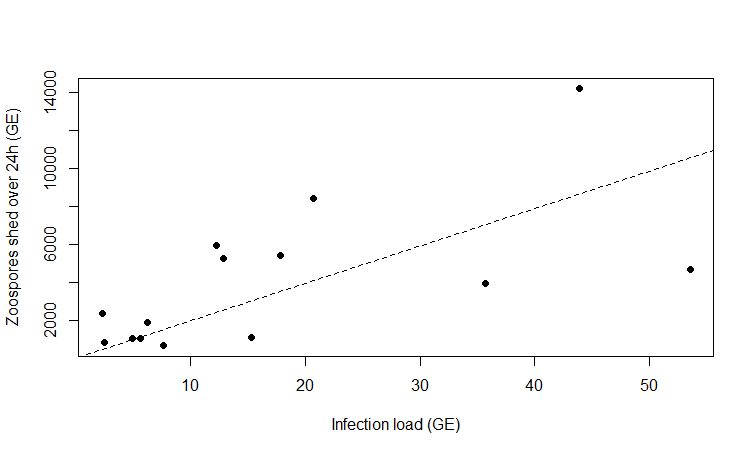

Fig S2. Relationship between on-host infection load (*L*, genomic equivalents; GE) and estimated number of zoospores shed over 24 h ($\lambda$), from data in Allen (2002). Dashed line shows fitted linear relationship $\lambda=196.17*L$ (adjusted R^2^ = 0.68).

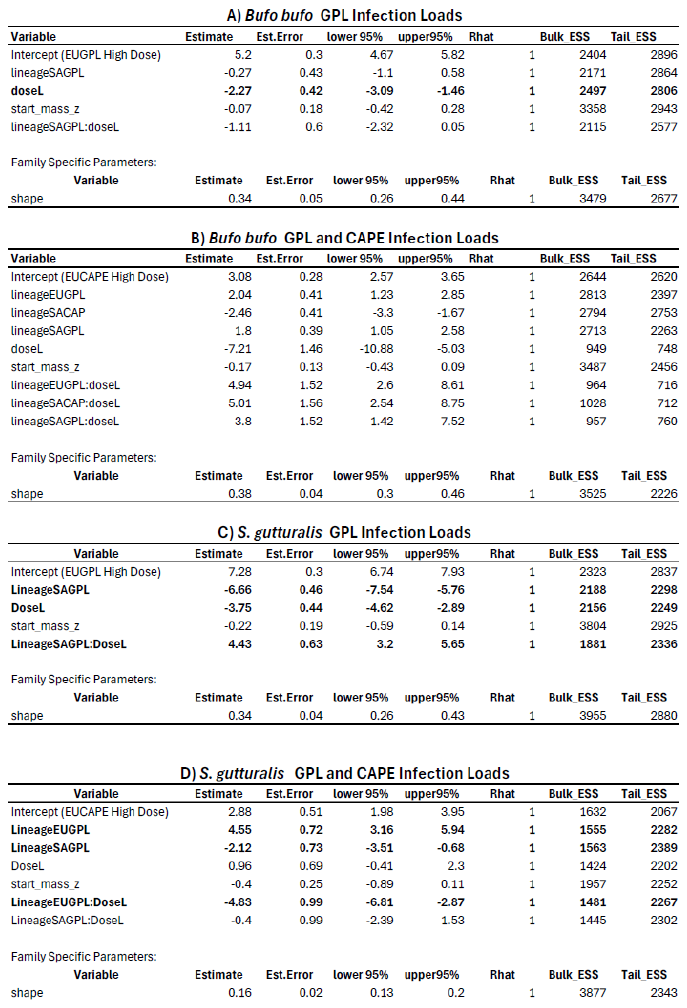

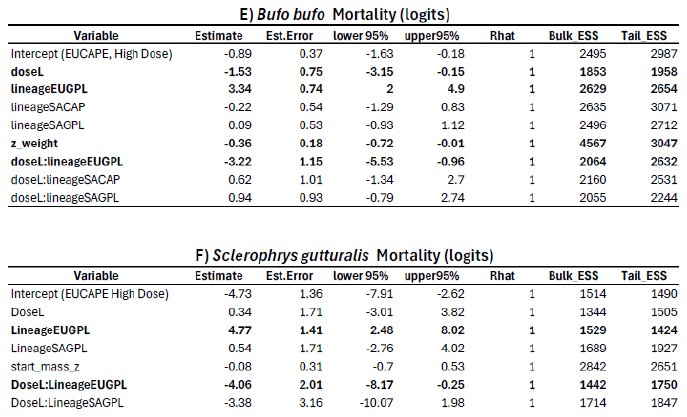

Table S1

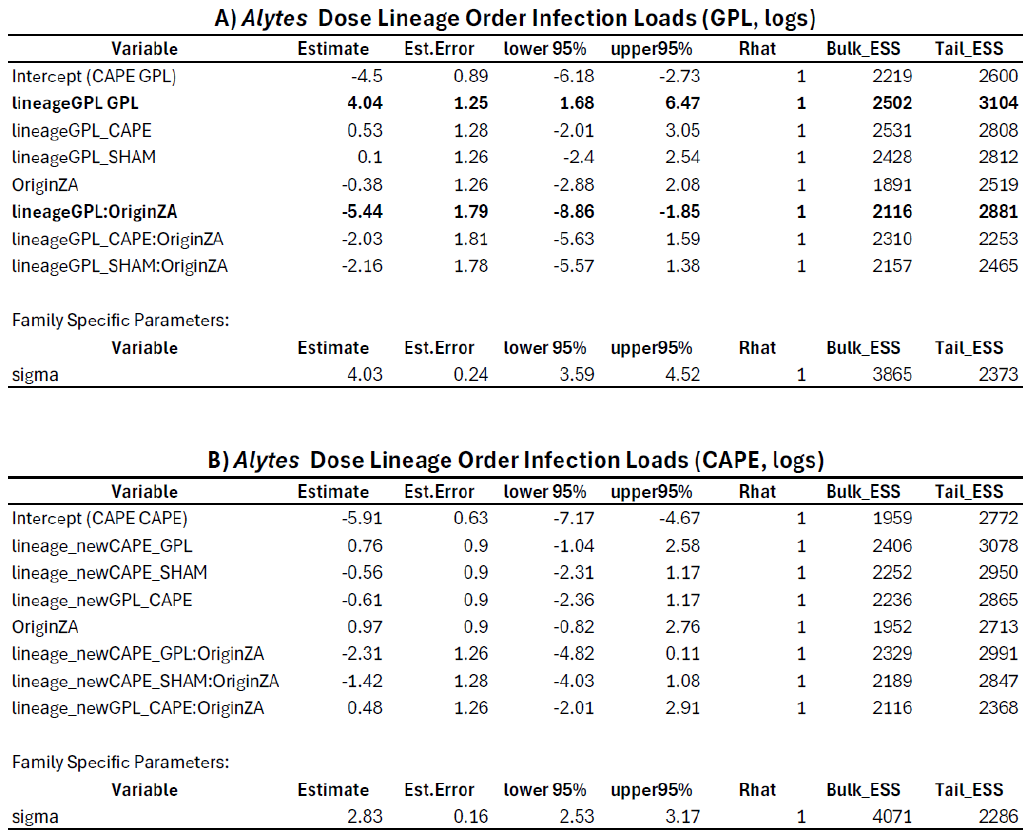

Table S2

| Focal lineage | Treatment | Number infected  (out of 20 exposed) | Probability of successful infection ($\sigma_{i}$) | Median infection load  (GE) | Zoospore shedding rate  ($\lambda_{i}$ day^-1^) |
| --- | --- | --- | --- | --- | --- |
| EU *Bd*GPL | Single | 5 | $\sigma_{UI}$ = 0.25 | 28.48 | 5587.9 |
| EU *Bd*GPL | Coexposure | 5 | $\sigma_{RI}$ = 0.25 | 43.56 | 8545.6 |
| EU *Bd*CAPE | Single | 1 | $\sigma_{UI}$ = 0.05 | 6.16 | 1208.9 |
| EU *Bd*CAPE | Coexposure | 1 | $\sigma_{RI}$ = 0.05 | 4.17 | 817.3 |
| ZA *Bd*GPL | Single | 4 | $\sigma_{UI}$ = 0.20 | 0.02 | 4.46 |
| ZA *Bd*GPL | Coexposure | 4 | $\sigma_{RI}$ = 0.20 | 31.63 | 6204.5 |
| ZA *Bd*CAPE | Single | 0 | $\sigma_{UI}$ = 0.001† | - | 1.96† |
| ZA *Bd*CAPE | Coexposure | 4 | $\sigma_{RI}$ = 0.20 | 16.06 | 3151.0 |

Table S3. Parameter estimates for the different lineages, under either single- or coexposure treatments. Data from low exposure treatments were used in all cases. The coexposure treatment involves hosts exposed to the focal lineage after prior exposure by the alternative lineage. Shedding rates were estimated from the infection load – shedding relationship calculated from data for *A. muletensis* in Allen (2022). †Although no infections were observed among the 20 exposed animals for this treatment, $\sigma_{UI}$ was set at an arbitrarily low value (1% prevalence), and an arbitrary median infection load of 0.01GE was assumed (resulting in estimated shedding rate of ~2 day^-1^), to allow for the possibility of very rare infections for this treatment.

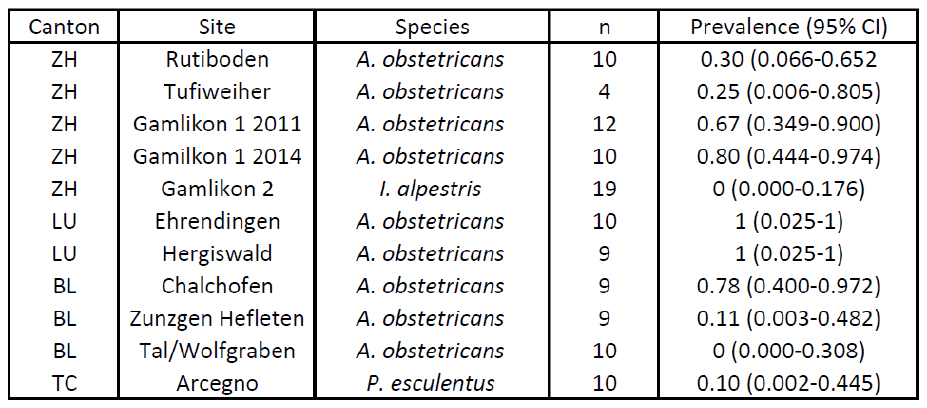

Table S4. Swiss Canton, region, host species, sample sizes and prevalence data for field surveys for *Bd*CH. *A. obstetricans* were tadpoles, all other species were adults. The complete data set is available from the Swiss National Biodiversity Data and Information Centre at https://doi.org/10.15468/ggwedn

ADDITIONAL REFERENCES

Bürkner P (2018) Advanced Bayesian Multilevel Modeling with the R Package brms. The R Journal, 10(1), 395-411. doi:10.32614/RJ-2018-017

Bürkner P (2021) Bayesian Item Response Modeling in R with brms and Stan. Journal of Statistical Software, 100(5), 1-54. doi:10.18637/jss.v100.i05

DiRenzo GV, Langhammer PF, Zamudio KR, Lips KR (2014). Fungal infection intensity and zoospore output of *Atelopus zeteki*, a potential acute chytrid supershedder. PLoS ONE, 9(3), e93356

R Core Team (2022). R: A language and environment for statistical computing. R Foundation for Statistical Computing,Vienna, Austria. URL <https://www.R-project.org/>.

Reeder NMM, Pessier AP, Vredenburg VT, Litvintseva AP (2012). A reservoir species for the emerging amphibian parasite Batrachochytrium dendrobatidis thrives in a landscape decimated by disease. PLoS ONE 7(3): e33567
